# Single-cell immune profiling with TCR clonotype barcoding identifies biomarker signatures that predict response to immune checkpoint blockade

**DOI:** 10.1101/2021.04.13.439713

**Authors:** Dean Lee, Ross Fulton, Monika Manne, Liang Schweizer, Andreas Raue

## Abstract

The identification of predictive biomarkers for patient treatment response is urgently needed to increase the probability of success of existing and novel experimental therapies. Single-cell profiling has provided novel biological insights into drug responses in the tumor microenvironment, but its potential for biomarker discovery has not been fully explored for therapeutic purposes. We describe a novel approach to discover predictive response biomarkers from single-cell data from a small patient cohort using the T cell receptor sequence intrinsic to each T cell to match clonotypes between pre- and post-treatment tumor samples. As a result, we have identified a predictive gene expression signature for immune checkpoint blockade and validated its predictive performance using data from three larger clinical studies. Our results demonstrated that applying clonotyping with single-cell genomic profiling is a promising novel approach for biomarker identification that does not require data collected from large patient cohorts. This could increase success rates, reduce clinical trial size, and significantly impact future clinical developments of immunomodulatory therapeutics.

## Introduction

Immune checkpoint blockade (ICB) has shown promising results in the clinic^1^, but only a portion of patients respond to the treatment. Various predictive biomarkers for treatment response such as tumor mutational burden^2,3^, microsatellite instability^4^, tumor T cell infiltration^5^, and PD-1 or PD-L1 status^6–8^ are used^9^; however, their ability to enrich for responder patients remains limited^10^. In addition, predictive gene expression signatures have been proposed using large-scale transcriptomic data collected from clinical trials^11^, although their effectiveness have only been shown retrospectively, after a large clinical study has been executed. The need for predictive biomarkers is even more pressing for therapies that are under development; due to a lack of such biomarkers, novel experimental therapies may fail to generate sufficiently broad clinical responses in early clinical trials and risk premature discontinuation before adequate data on responding patients can be acquired^12–14^. Therefore, the identification of predictive biomarkers, especially from small patient cohorts, is urgently needed to maximize the chances of success of novel experimental therapies.

The discovery of predictive biomarkers, with ICB as a case study here, typically relies either on hypotheses informed by the drug mechanism^15,16^ or on the retroactive analysis of bulk or single-cell RNA-seq data^17^. The former risks missing relevant factors that are outside the narrow set of starting hypotheses, while the latter often leads to complex signatures that can be hard to interpret^18,19^. Furthermore, bulk RNA-seq analysis cannot differentiate between individual cell states and changes in cell frequencies, which confounds the resulting signatures. More recently, single-cell profiling studies have provided novel biological insights into drug responses in the tumor microenvironment (TME), but its potential for biomarker discovery has not yet been fully leveraged. Some of these studies have investigated the dynamics of T cell clonality highlighting the importance of both preexisting and newly infiltrating T cell clones and their expansion^18,20^. Based on these findings, we hypothesized that T cells, especially those with known cytotoxic roles, that expand after ICB in patients are more likely to contain predictive biomarkers for a positive clinical outcome. Given that some T cells subsets, such as Tregs, can have immune-suppressive effects, we further hypothesize that the lack of expansion of these cells could also lead to a positive clinical outcome.

Using published single-cell RNA-sequencing^20^, we describe a novel approach to discover predictive biomarkers that uses T cell receptor (TCR) clonotyping as a barcoding strategy across pre- and post-treatment tumor samples. By linking T cells between pre- and post-treatment tumor samples via their clonotype, this approach can identify predictive gene expression signatures from cells in the pre-treatment samples that distinguish responsive from non-responsive cells. Using this clonotype barcoding (CB) approach, we identified a predictive gene expression signature from a small patient cohort of ICB treated patients and validated its predictive performance using data from three larger ICB clinical studies.

## Results

### Clonal expansion of T cells after ICB

To investigate which T cells responded to ICB by clonal expansion, we used data from 14 skin cancer patients each with site-matched tumor biopsies from pre- and post-anti-PD-1 treatment. Each biopsy was profiled with single-cell RNA-sequencing (scRNA-seq) for gene expression and single-cell TCR-sequencing (scTCR-seq) for clonotype analysis^20^. We separated the samples from basal cell carcinoma (BCC) and squamous cell carcinoma (SCC) for further analysis. We then used clustering and dimensionality reduction algorithms to identify cell populations that share common features based on their gene expression profiles and found clusters that separated by both T cell subset (**Fig. 1a, b**) and patient-specific identifiers (**Fig. 1c, d**). Clonal expansion triggered by ICB was quantified by identifying T cell clonotypes which match between the pre- and post-treatment samples to calculate the treatment-induced fold-increase of the clonotype size (**Fig. 1e, f**). In the BCC data, 2.02% of all CD8+ T cells, 0.63% of all conventional CD4+ T cells, and 0.99% of all FOXP3+ CD4+ Tregs expanded after ICB, with an average fold-increase of 3.82, 2,75, and 4.54, respectively. While CD8+ T cells predominantly responded in BCC, in SCC there was a broader T cell response with 5.14% of all CD8+ T cells, 5.04% of all CD4+ T cells, and 2.70% of all Tregs expanding after ICB, with an average fold-increase of 4.88, 5.26, and 4.06, respectively. The absolute number of Treg clonotypes that responded to ICB with clonal expansion was limited in both datasets. Therefore, we combined Tregs and conventional CD4+ T cells for subsequent analysis. To increase resolution, we repeated the clonotype expansion analysis on each major T cell subset and found that in BCC, the majority of responding CD8+ T cells came from one patient (**Suppl. Fig. 1**). However, still only 6.74% of single CD8+ T cells of this patient were identified as responding. In the SCC data, the responding cells were more dispersed across patients. In conclusion, we were able to identify both CD8+ and CD4+ T cell clonotypes that expanded in response to ICB based on scTCR-seq clonotype profiling in BCC and SCC cohorts.

**Fig. 1.**
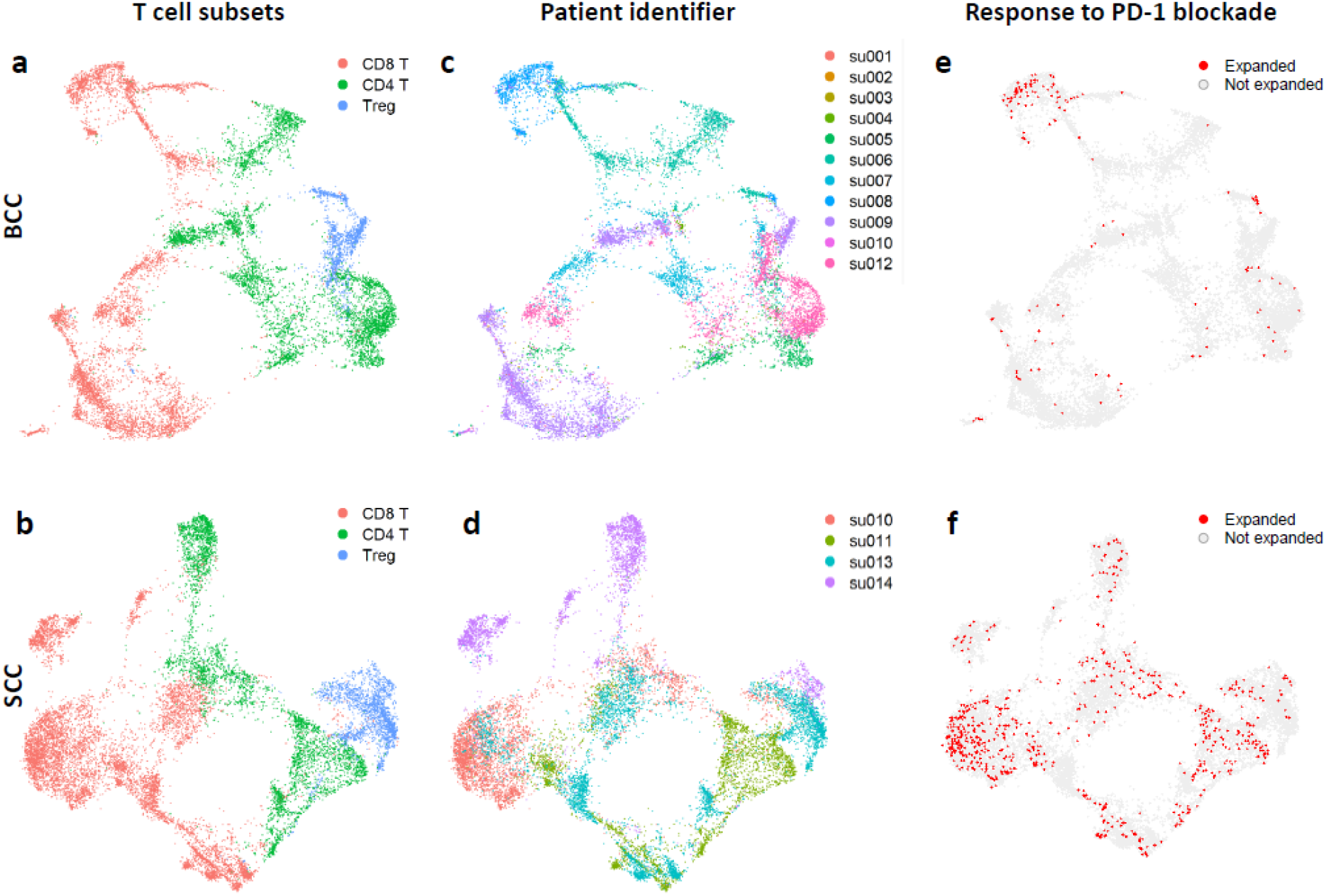
Classification of expanded and non-expanded T cell subsets from patients treated with ICB. **a-f**, 12,551 T cells from BCC (**a, c, e**) and 14,522 T cells from SCC (**b, d, f**) were clustered and visualized with dimension reduction algorithms based on the similarity of their gene expression profiles. Association of single-cells with CD8+, CD4+ or regulatory T cell subsets was identified based on marker gene expression (**a, b**), and association of single-cells with patient identifiers is indicated (**c, d**). T cells that expanded in response to anti-PD-L1 treatment were identified based on scTCR-seq data and are marked in red. All other T cells are displayed in gray (**e, f**).

### Differentially expressed genes in pre-treatment T cell clonotypes that expanded after ICB

Next, we investigated if we could use *pre-treatment* gene expression profiles to distinguish T cells that responded to ICB with clonal expansion from T cells that did not respond. Using our CB approach, we grouped the TCR clonotypes in the pre-treatment samples into a fraction that did respond and a fraction that did not respond after ICB (Suppl. Fig. 1). We then identified genes that were differentially expressed in the set of responding cells using their scRNA-seq gene expression profiles. To identify differentially expressed genes, we calculated the gene expression fold-change between the two fractions for each gene and used several filtering steps to ensure the identified genes represent clear (not affected by sparsity of single-cell data or high baseline expression), meaningful (have an appreciable log fold-change), and statistically robust signals (significant based on multiple-testing-corrected rank-sum test) (**Suppl. Fig. 2**). In BCC patient samples, we found 8 genes (*IFNG, ADGRE2, TNF, FGL2, RHOB, ADRB2, NFKBIZ, STAT1*) that were upregulated and 9 genes (*TRAT1, MRPL18, PEBP1. CMTM7, CCR7, CYSTM1, AHSA1, CMSS1*) that were downregulated in CD8+ T cells relative to clonotypes that did not expand following ICB (**Fig. 2a**). In contrast, there were no differentially expressed genes in CD4+ T cells (**Fig. 2b**). This may be due to the low frequency of expanding cells in this subset (0.7%). In SCC samples we did not find any differentially expressed genes in CD8+ T cells (**Fig. 2c**), but we identified 5 downregulated (*LAIR2, ACP5, IL1R2, GZMA, FOXP3*) and two upregulated (*G0S2, BAG3*) genes in CD4+ T cells relative to clonotypes that did not expand following ICB (**Fig. 2d**). In summary, we identified two distinct sets of differentially expressed genes at the single-cell level, one linked to CD8+ T cells in BCC and another linked to CD4+ T cells in SCC, that correlated with the clonal expansion in response to ICB.

**Fig. 2.**
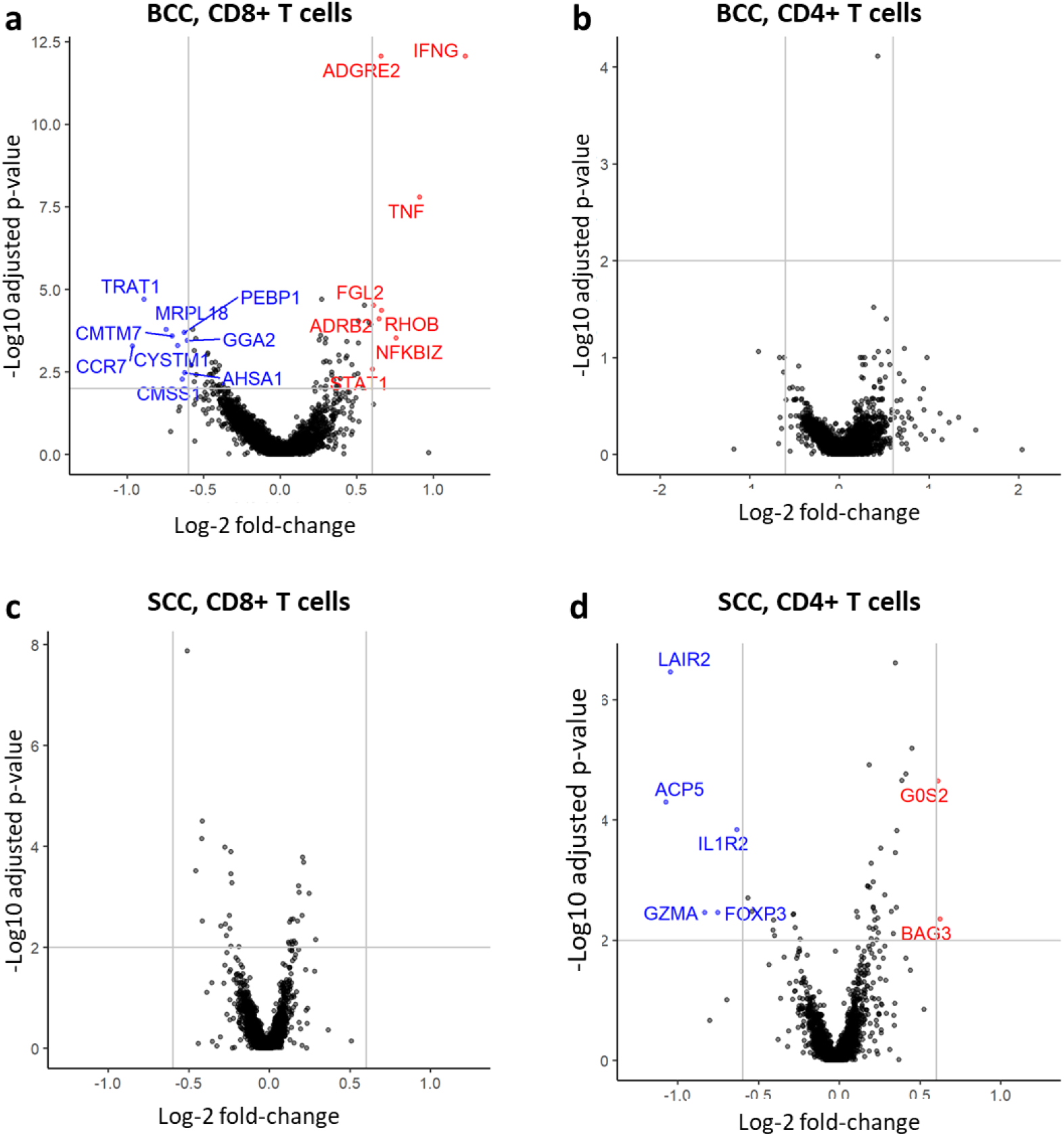
Differentially expressed genes that distinguish T cells that will expand after PD-1 blockade based on their pretreatment gene expression profile. **a-d**, Samples are divided into BCC (**a, b**) and SCC (**c, d**), and CD8+ (**a, c**) and CD4+ T cells (**b, d**). To select for the most clearly defined differentially expressed genes, we removed genes that have a high background expression, pass threshold for both fold-change and statistical significance (gray lines). The resulting list of differentially expressed genes are named and highlighted in color, where blue means downregulation on the expanded fraction, and red mean upregulation in the expanded fraction.

### Generation of predictive signatures of T cell expansion from scRNA-seq data

Given the sets of differentially expressed genes in CD4+ and CD8+ T cells, we wanted to find which single-cell gene expression signature best predicted the response to ICB that could later be applied to bulk gene expression data to predict patient response to ICB in clinical trials. We noted that bulk RNA-seq data not only represents tumor-infiltrating T cells, but the mixture of all cells in the TME, including immune, stromal, and malignant cells. To obtain a T cell-specific predictive signature, we restricted the list of differentially expressed genes to genes with expression more restricted to T cells subsets by computing their correlations with T cell lineage markers. To obtain a gene expression signature that best predicted T cell expansion after ICB at the single-cell level, we trained a logistic regression model on the remaining differentially expressed genes (**Suppl. Fig. 2**). We further investigated whether more stringent cut-offs for differential expression, which lead to decreased numbers of genes in the signature, would impact the predictive performance of the logistic regression models (**Tab. 1**). For the CD8+ T cell expansion signature, we found that the most parsimonious model with just one gene, *IFNG*, had the highest accuracy and specificity as a positive predictor of expansion (**Fig. 3a**, c.f. Suppl. Fig. 1e). For the CD4+ T cell signature, we found that a model with two genes, *GZMA* and *FOXP3*, had the best accuracy and specificity (**Fig. 3b**, c.f. Suppl. Fig. 1h); CD4+ T cell transcription of both these genes were *negative* predictors of expansion (i.e. prior to treatment these genes were downregulated in clonotypes that did respond versus those that did not respond after ICB). This signature applies to both Tregs and a fraction of conventional CD4+ T cells. In conclusion, we identified gene expression signatures that predicted response to ICB at the single-cell level: expression of one signature was a positive predictor of CD8+ T cell expansion (CD8-expansion signature) and, inversely, expression of the other signature was a negative predictor of CD4+ T cell expansion (CD4-non-expansion signature).

**Tab. 1.**
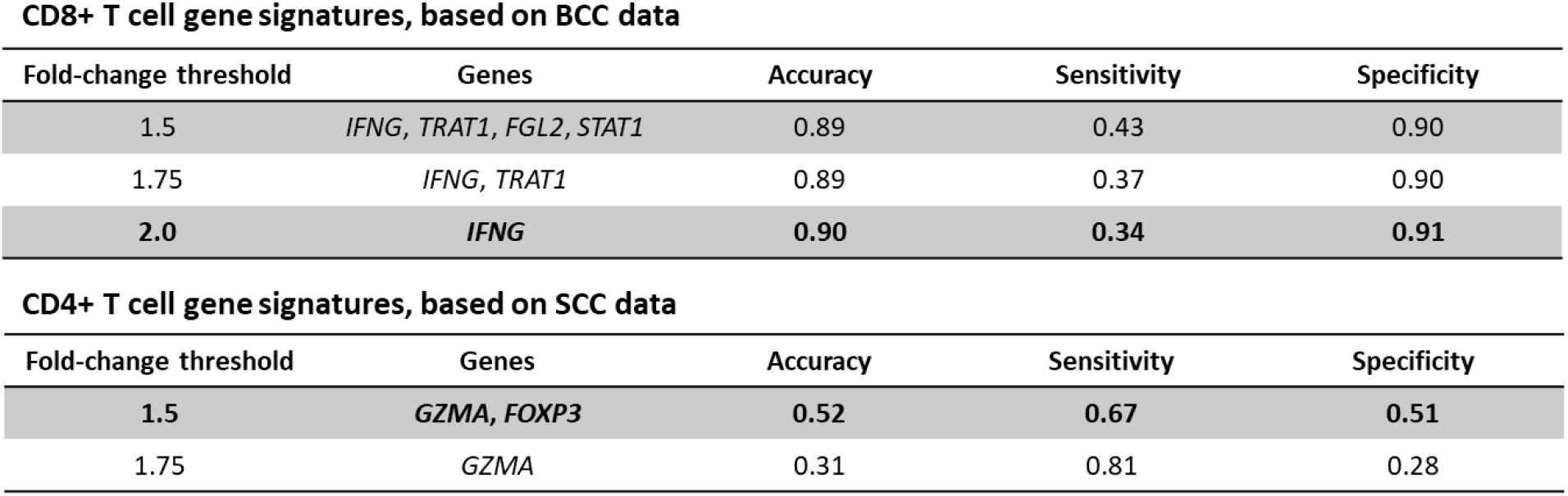
Results from logistic regression model training for deriving gene expression signatures that can optimally predict T cell expansion after PD-1 blockade. The table show the results from training logistic regression models to predict the T cell expansion after PD-1 blockade at the single-cell level. For each of the T cell subsets independent signature have been derived, based on the analysis differential gene expression. For each subset, the threshold for differential gene expression fold-change was varied to investigated whether more stringent cut-offs would impact the predictive performance. The predictive performance was measured by accuracy (proportion of correct predictions, both true positives and true negatives, among the total number of cases), sensitivity (true positive rate), and specificity (true negative rate).

| **CD8+ T cell gene signatures, based on BCC data** |  |  |  |  |
| --- | --- | --- | --- | --- |
| Fold-change threshold | Genes | Accuracy | Sensitivity | Specificity |
| 1.5 | *IFNG, TRAT1, FGL2, STAT1* | 0.89 | 0.43 | 0.90 |
| 1.75 | *IFNG, TRAT1* | 0.89 | 0.37 | 0.90 |
| 2.0 | *IFNG* | 0.90 | 0.34 | 0.91 |
| **CD4+ T cell gene signatures, based on SCC data** |  |  |  |  |
| Fold-change threshold | Genes | Accuracy | Sensitivity | Specificity |
| 1.5 | *GZMA, FOXP3* | 0.52 | 0.67 | 0.51 |
| 1.75 | *GZMA* | 0.31 | 0.81 | 0.28 |

**Fig. 3.**
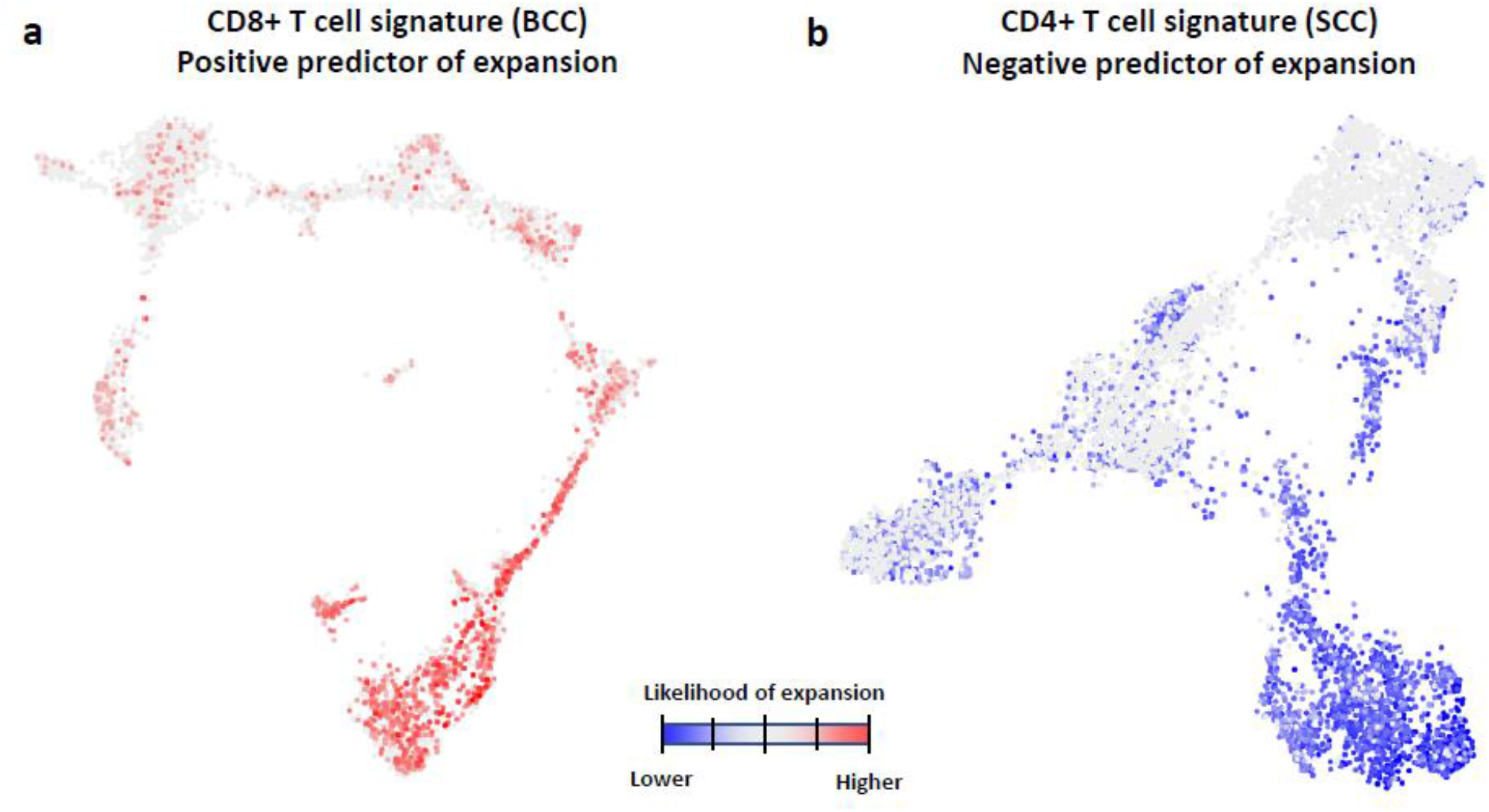
Illustration of the score values of the identified predictive gene expression signatures as overlay on the single-cell data visualization. **a, b**, Two distinct signatures were identified, a CD8+ T cell signature in BCC patient samples that was a positive predictor of expansion (**a**), and a CD4+ T cell signature in SCC patient samples that was negative predictor of expansion (**b**). Signature score values are relative, unitless and indicated by a color scale with the score values shown in red that correlated with high likelihood of expansion and blue color indicating lower likelihood of expanding.

### Predictive signatures of T cell expansion can stratify patients in clinical trial data

To investigate if the two gene expression signatures derived from single-cell data (**Fig. 3)** can predict ICB treatment outcome, we applied both the CD8-expansion signature and the CD4-non-expansion signature to data from three phase II clinical trials with an anti-PD-L1 checkpoint inhibitor (atezolizumab) where bulk RNA-seq on pre-treatment patient samples was available: the POPLAR study^16^ (NCT01903993) in locally advanced or metastatic non-small-cell lung cancer, the IMvigor210 study^21^ (NCT02951767, NCT02108652) in locally advanced or metastatic urothelial bladder cancer, and the IMmotion150 study^22^ (NCT01984242) in advanced renal cell carcinoma. For each study, we generated a signature score for each patient, used a median cutoff to distinguish signature-high from signature-low patients, and investigated if the signature can predict outcome to treatment using survival analysis (**Suppl. Fig. 3**). For the CD8-expansion signature, we found that signature-high patients trended toward longer progression-free survival (PFS) (**Fig. 4a**). For the CD4-non-expansion signature, we found that signature-high patients also tended to experience longer PFS (**Fig. 4b**). This was surprising because a high signature means that CD4+ T cells in their tumors would have a lower probability to expand post ICB. We found expression of *FOXP3* and *GZMA*, the genes contributing to this signature, within distinct conventional CD4+ T cell clusters, however, there was also some overlap in the Treg cluster (**Suppl. Fig. 4**). Therefore, we cannot exclude the potential predictive importance of this nonexpansion signature originating from within the Treg subset. While both signatures significantly (p-value < 0.05) enriched for responding patients in IMvigor210 and POPLAR, only patients in the IMmotion150 trial with the CD4-nonexpansion signature reached significance in the combination arm (atezolizumab + bevacizumab). Next, we asked if applying both signatures in combination can further improve treatment outcome predictions. We divided the patients into four groups with the CD8 signature indicated first: above median score for both signatures (Pos-Pos), above median score for one signature (Pos-Neg, Neg-Pos), or below median score for both signatures (Neg-Neg). We found that a Pos-Pos combined signature could further improve the predictions for IMvigor210 and the combination arm of IMmotion150 (**Fig. 4c**). For POPLAR, we found that all groups positive for at least one of the two signatures had significantly improved outcome relative to the low-low group (**Fig. 4c**). Based on these observations, we derived a combined signature that takes both the signatures into account. When compared with other published signatures and gene expression markers of interest including *CD274* (PD-L1), *PDCD1* (PD-1), *CD8A, CD4*, and *FOXP3*, we found that our combined signature consistently achieved similar or superior hazard ratios compared to other signatures or markers of interest (**Suppl. Tab. 1**). Of note, for the IMmotion150 atezolizumab monotherapy arm, none of the signatures was able to provide significantly lowered hazard-ratios (HRs). In summary, we demonstrated that gene expression signatures derived from pre-treatment scRNA-seq data from a small number of patients can predict outcome in larger cohorts of ICB treated patients when applied to pre-treatment bulk RNA-seq data. Furthermore, we showed that a combination signature can lead to further improved predictions depending on the tumor type.

**Fig. 4.**
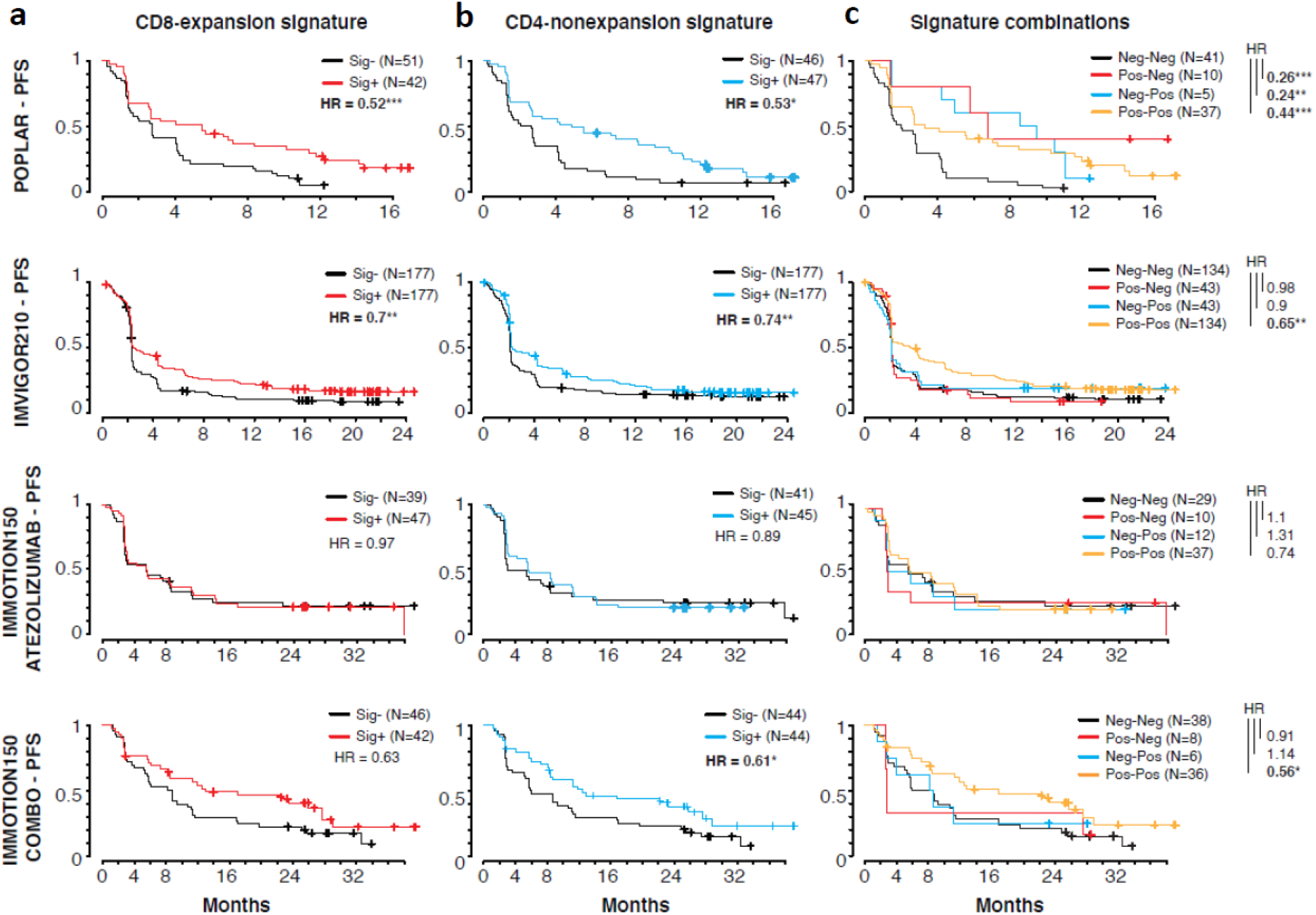
Survival analysis of clinical trial data using the predictive signatures. **a**, Kaplan-Meier plots of progression-free survival (PFS) are shown for the atezolizumab treated patients in each clinical trial, with patients divided by their predictive signature scores above (Sig+) or below (Sig-) the median among all patients in the corresponding clinical trial. Censored observations are indicated by a plus symbol. Hazard ratios from a Cox proportional-hazard model analysis are shown (* p_val_ < 0.05, ** p_val_ < 0.01, *** p_val_ < 0.001). **b**, same as in **a**, except that patients were divided into four groups (low-low, low-high, high-low, and high-high) based on the two signatures shown in **a**, with the CD8 signature indicated first. Hazard ratios and significance relative to low-low groups are indicated.

### Outcome prediction in overall population and comparison to previously published signatures

To further demonstrate how predictive signatures derived from single-cell data could impact the clinical development of novel experimental drugs, we analyzed how patient stratification using our combination signature could improve the probability of obtaining a significant improved PFS in a clinical trial. For this we used a subset of patient data of the POPLAR and IMmotion150 studies in which pre-treatment bulk RNA-seq data were available for the experimental and standard-of-care comparator arms. We applied our combination signature to divide patients treated with ICB in each trial into biomarker-positive (BM+) and biomarker-negative (BM-) groups. For each trial and biomarker group, we then performed a survival analysis between the comparator arms and the experimental arms and compared the results to the outcome in the unstratified patient population (**Tab. 2**). In the unstratified case, the experimental arms did not significantly improve HRs in any of the three analyses. After stratifying patients based on our combination signature, we found overall lowered HRs in the BM+ groups as compared to unstratified. Statistical significance was reached for the IMmotion150 trial combination arm, suggesting that the combination of atezolizumab and bevacizumab is more beneficial than the multi-targeted receptor tyrosine kinase (RTK) inhibitor sunitinib in this patient population. Conversely, in the BM-groups we found increased HRs as compared to unstratified. Significance was reached in the BM- group of POPLAR, suggesting that atezolizumab is less beneficial than the chemotherapy docetaxel in this patient group. Correspondingly, in the BM+ group of POPLAR we found a strong trend towards decreased HR as compared to unstratified; however, it did not formally reach significance (p = 0.06). We also compared our combination signature to previously published signatures and other biomarkers of interest (**Tab. 2**). We found that our signature consistently showed better or similar performance to other markers of relevance. Of note, for the IMmotion150 atezolizumab monotherapy arm, none of the signatures was able to provide significant lowered HRs. In summary, we demonstrated that gene expression signatures derived from single-cell data from a small number of patients could prospectively identify populations of patients that have a greater chance of benefiting from an experimental treatment compared to patients in an unstratified patient population.

**Tab. 2.** Comparison of hazard-ratios between experimental and comparator arms in both biomarker positive (BM+) and negative (BM-) populations. Hazard ratios and two-sided statistical test from a Cox proportional-hazards model on patients in both groups are shown with significance indicated (* p_val_ < 0.05, ** p_val_ < 0.01, *** p_val_ < 0.001). ^†^p_val_ of 0.06.

| Stratification strategy | POPLAR<br>Atezolizumab vs. Docetaxel |  | IMmotion150<br>Atezolizumab vs. Sunitinib |  | IMmotion150<br>Atezolizumab + Bevacizumab vs. Sunitinib |  |
| --- | --- | --- | --- | --- | --- | --- |
| Unstratified | 0.93 |  | 1.08 |  | 0.81 |  |
|  | *BM+* | *BM-* | *BM+* | *BM-* | *BM+* | *BM-* |
| Expansion signature | 0.68<sup>†</sup> | 1.79* | 0.9 | 1.35 | 0.56* | 1.14 |
| Wu et al.<sup>18</sup> | 0.87 | 1.03 | 0.88 | 1.39 | 0.55* | 1.23 |
| CD274 (PD-L1) | 0.8 | 1.06 | 0.9 | 1.4 | 0.68 | 0.98 |
| PDCD1 (PD-1) | 0.64* | 1.39 | 0.83 | 1.44 | 0.48** | 1.4 |
| CCL5 | 0.74 | 1.27 | 0.75 | 1.53 | 0.52** | 1.33 |
| CD8A | 0.74 | 1.2 | 0.86 | 1.39 | 0.51** | 1.35 |
| CD4 | 1 | 0.87 | 1.1 | 1.09 | 0.72 | 0.96 |
| FOXP3 | 0.74 | 1.2 | 0.95 | 1.27 | 0.63 | 1.06 |

## Discussion

Using pre- and post-treatment single-cell data from patients who received ICB, we have demonstrated a novel approach to identify predictive gene expression signatures as biomarkers that stratify patients to improve the outcome of clinical trials. Our clonotype barcoding (CB) approach uses the TCR sequence intrinsic to each T cell to match clonotypes between pre- and post-treatment tumor samples and to identify T cells that increased in clonotype size in post-treatment samples. While this increase could be attributed to clonotype expansion of pre-existing clones, we have not explored the relative contribution of better retention or higher recruitment. The CB approach prioritizes gene expression patterns from expanded and recurring T cell clonotypes detected in the TME for predicting treatment outcome.

Single-cell profiling enables the CB approach to exploit heterogeneity at the single-cell level to derive predictive gene expression signatures. In contrast to conventional approaches that require larger numbers of patient samples to compare different groups of patients, linking groups of T cells from pre- and post-treatment samples from the same patients provides an elegant way to identify the most critical predictive parameters within a smaller patient cohort, such as those available from an early clinical study. The signatures we found using the CB approach were surprisingly simple and amenable to interpretation and practical clinical implementation (i.e., immunohistochemistry or *in situ* hybridization) but would have been missed otherwise as part of more complex, retroactively defined signatures.

The single-cell data used in this study, though limited, was unique at the time of this analysis in that it included both pre- and post-treatment samples from the same patients receiving ICB. While it is still limited in the number of matching T cell clonotypes pre- and post-treatment and in the number of patients exhibiting strong clonotype size increases, we anticipate that the generation of scRNA-seq datasets with larger numbers of patient samples and profiled cells will yield more refined signatures that more clearly connects T cell biology to patient responses to immunotherapies. Furthermore, better optimized, and standardized single cell data generation and analysis will enrich the data set further and provide more meaningful analysis.

Mechanistically, the identification of *IFNG* as the best predictor of CD8+ T cell clonal expansion in BCC patients treated with ICB indicates the presence of functional tumor-reactive cells actively recognizing cognate tumor antigens but whose expansion was inhibited by the PD-1/PD-L1 pathway^11,16,23^. It was therefore surprising that a CD8+ T cell signature in SCC patients was not identified, in spite of a greater proportion of pre-treatment CD8+ T cell clones expanding (2.02% and 5.14% of CD8+ T cell clones in BCC and SCC, respectively). T cell clonal responses in SCC were broader than in BCC, with a greater proportion of conventional CD4+ T cells and Tregs expanding in SCC, which may indicate distinct ICB mechanisms of action between these cancer indications. Whereas CD8+ T cell expansion in BCC may have been a direct consequence of ICB, it may have been an indirect consequence in SCC, potentially resulting from CD4+ T cell help^24,25^, decreased regulatory T cell inhibition^26,27^, or from anti-tumor activity from immune cell subsets not captured in our CB approach, such as NK cells^28,29^.

Interpretation of the CD4-nonexpansion signature is more complex. *FOXP3* and *GZMA* do not always correspond to the same cell subset (**Suppl. Fig. 4**), and *GZMA* by itself still had predictive power outside of the Treg cluster. While the transcription factor FOXP3 is characteristic of Tregs, *FOXP3* transcription is also expressed in conventional T cells following activation^30^, and indeed *FOXP3* was found within the conventional CD4+ T cell cluster in the SCC scRNA-seq data. *GZMA*, which encodes the cytolytic protein granzyme A and is associated with effector T cells, has been previously identified as a positive prognosticator^31^ or a predictive biomarker for ICB treatment^32^, although this is typically attributed to CD8+ T cells. However, since there is some overlap between *FOXP3* and *GZMA* within the Treg cluster (**Suppl. Fig. 4**), we cannot exclude the potential predictive importance of this Treg subset. Granzyme A produced by activated Tregs reportedly makes these cells more susceptible to self-inflicted apoptosis^33^, but this will require further exploration.

Further, we demonstrated that our predictive gene expression signatures identified using the CB approach can be validated in larger clinical trials. We showed that both the CD8-expansion signature as well as the CD4-non-expansion signature can predict favorable outcomes to treatment and that a combination of both signatures can lead to further improved results. We additionally note that the predictive signatures, though derived based on data from skin cancer patients, still showed predictiveness in larger cohorts of three other solid cancer types. However, the best combination of signatures depended on the treated patient population.

Finally, we demonstrated how the predictive signatures could be applied for patient stratification to improve the probability of success of clinical trials. While our signatures consistently performed better or similarly to other relevant markers, we noted that PD-1 expression alone was also a strong predictive biomarker. This is not unexpected based on our mechanistic understanding of ICB^6–8^. Importantly, the strength of our CB approach was that it could identify a biomarker signature in an unbiased way that performed similarly well. This could be most relevant for identifying biomarkers for novel experimental therapies where the mechanism may be less understood or more complex. We conclude that our proposed CB approach is a promising novel use of single-cell data for biomarker identification that could increase the success rates of future clinical trials for novel immunomodulatory therapeutics by enriching for responders and reducing trial size.

## Materials and Methods

### Identification of clonal expansion based on scTCR-seq data

TCR clonotype labels for individual T cells were obtained from Yost et al.^20^. Clonotypes shared by pre- and post-treatment T cells within the same patient were identified based on their matching TCR CDR3 sequences. Clonotypes with post-treatment clonotype size (number of T cells sharing the same TCR CDR3 sequences) greater than the pre-treatment clonotype size were considered to have expanded in response to ICB. For patient su013 in the SCC dataset, 4488 T cells were profiled in the pre-treatment sample but only 69 in the post-treatment sample; this imbalance in T cell counts partially explains why no clonotypes in patient su013 were considered to have experienced expansion after treatment. Patient su010 presented with both BCC and SCC lesions; therefore, this patient was represented in both the BCC and SCC datasets.

### Clustering and dimensionality reduction of scRNA-seq data

Clustering of scRNA-seq data from all BCC cell types was performed as previously described^20^ with Seurat (version 3.1.1). Based on this first round of clustering and the expression of canonical T cell markers (*CD8A*, *CD4, FOXP3*), we identified T cell clusters. The T cells specifically from pre-treatment samples were extracted from the complete dataset then re-clustered. In this round of clustering, the default variable gene selection criteria in Seurat was used. These variable genes were used for principal component analysis (PCA). The first 20 principal components were used for shared nearest neighbor-based clustering with k = 20 and resolution = 3. The same principal components were used for visualization with UMAP projections with minimum distance = 0.1 and number of neighbors = 20. The SCC scRNA-seq dataset^20^ was already specific to T cells. After extracting the pre-treatment T cells only from this dataset, clustering was performed with Seurat. These pre-treatment T cells were clustered the same way as BCC pre-treatment T cells. After the pre-treatment T cells were clustered, each cluster was annotated as CD8+ T cells, conventional CD4+ T cells, or Tregs based on expression of *CD8A, CD4*, or *FOXP3*, respectively.

### Differential gene expression analysis

Using only pre-treatment scRNA-seq data, differential gene expression analysis was performed to determine significantly up- or down-regulated genes in the expanded T cell population versus the non-expanded T cell population. Several filtering steps were taken to remove genes unlikely to be biologically relevant from testing. Testing was limited to genes that were detected in at least 10% of cells in either of the two populations being compared. Non-coding genes, as defined by HGNC, were removed from testing. Human T cell receptor alpha variable (*TRAV*) genes, human T cell receptor beta variable (*TRBV*) genes, and human leukocyte antigen (HLA) genes were also removed from testing. For the genes that passed filtering, the Wilcoxon Rank-Sum Test was performed to detect differential gene expression between the expanded T cell population versus the non-expanded T cell population. The Benjamini-Hochberg correction was used to adjust the resulting p-values to control the false discovery rate. Differentially expressed genes were defined to be those with a log2-fold-change > 0.6 and an adjusted p-value < 0.01. In addition, for genes to be considered significantly upregulated in the expanded T cell population, they were required to be expressed in less than 30% of the non-expanded T cell population. Likewise, for genes to be considered significantly downregulated in the expanded T cell population, they were required to be expressed in less than 30% of the expanded T cell population.

### Generation of predictive signatures based on differentially expressed genes

Differential gene expression analyses yielded two lists of differentially expressed genes: one from the BCC CD8+ T cell dataset and the other from the SCC CD4+ T cell dataset. For each list, genes with a fold-change >1.5 were used as predictors in a logistic regression classifier that predicted the expansion status of each cell. The classifier computed a coefficient for each gene that reflected how strongly the gene expression level of each gene predicted the expansion status of each cell (expanded vs. non-expanded). A positive coefficient indicated that a cell expressing that gene was more likely to be expanded. A negative coefficient indicated that a cell expressing that gene was more likely to be non-expanded. Then the process was repeated with a fold-change threshold of 1.75, then again with a fold-change threshold of 2.0; with increasing fold-change thresholds, the number of genes being used in the logistic regression classifier decreased. The coefficients, accuracy, sensitivity, and specificity of each logistic regression classifier at each fold-change threshold was recorded. For the BCC CD8+ T cell dataset, we compared the performance of the logistic regression classifiers and chose the most parsimonious model that did not suffer from a substantial drop in accuracy, sensitivity, and specificity. The top-performing classifier in this case contained one gene, *IFNG*, which has a positive coefficient. This was defined as the CD8-expansion signature. Using the same classifier selection criteria for the SCC CD4+ T cell dataset, we defined the CD4-nonexpansion signature as containing two genes, *GZMA* and *FOXP3*, each having a negative coefficient. For each gene in an expansion signature, the absolute value of its logistic regression coefficient was taken to be its weight.

### Prediction of patient response to ICB by application of single-cell-based gene expression signatures to bulk RNA-seq data

Bulk tumor RNA-seq data from three ICB clinical trials were used to assess the effectiveness of the CD8-expansion signature, the CD4-nonexpansion signature, and those two signatures combined in predicting patient response. In POPLAR (NCT01903993)^16^, patients with locally advanced or metastatic non-small-cell lung cancer were treated with atezolizumab (n = 93) or docetaxel (n = 100). In IMvigor210 (NCT02951767, NCT02108652)^21^, patients with locally advanced or metastatic urothelial bladder cancer (n = 354) were treated with atezolizumab. In IMmotion150 (NCT01984242)^22^, patients with advanced renal cell carcinoma were treated with atezolizumab alone (n = 86) or with both atezolizumab and bevacizumab (n = 88). For patients in each clinical trial, the weighted sums of the genes in a signature (CD8+ T cell expansion signature or CD4+ T cell expansion signature) were calculated to be the signature score. The median signature score was determined for each clinical trial, and patients who scored above that median were placed in the biomarker-positive group, while the remaining patients in the same trial were placed in the biomarker-negative group. Survival analyses comparing the biomarker-positive and the biomarker-negative groups in each clinical trial arm were performed as previously described by Wu et al^13^. A Cox proportional-hazards model was fitted to the survival data in each arm of each clinical trial; the resulting hazard ratio and p-value of the Wald statistic were reported. The results of these survival analyses were represented in Kaplan-Meier plots. To study the interaction between the CD8+ and CD4+ T cell expansion signatures within a clinical trial, we assigned the patients in each arm to one of four groups: above median score for both signatures (positive-positive), above median score for one signature (positive-negative, negative-positive), or below median score for both signatures (negative-negative). Survival analysis and Kaplan-Meier plotting were performed as described above, with the negative-negative group serving as the baseline comparator to each of the other three groups. As references, *CD274* (PD-L1), *PDCD1* (PD-1), *CD8A, CD4* and *FOXP3* were each treated as its own individual unweighted biomarker. Survival analysis comparing the biomarker-positive and the biomarker-negative groups in each arm of each clinical trial was performed as described above.

### Prediction of trial outcome (experimental vs. comparator arms) via predictive signatures

To compare the effectiveness of these T cell expansion signatures between the experimental (atezolizumab, atezolizumab + bevacizumab) and comparator (sunitinib) arms in IMmotion150, the CD8+ and CD4+ T cell expansion signatures were merged into a single signature, with the respective weights of each gene carrying over to the merged signature. First, to establish a baseline, survival analysis comparing each experimental arm with the comparator arm was done to detect any pre-stratification differences in survival. Next, the patients from IMmotion150 were stratified into a biomarker-positive group and a biomarker-negative group based on the merged signature. For each of the biomarker-stratified groups, survival analysis comparing atezolizumab-treated vs docetaxel-treated patients was performed as described above. For POPLAR, the same process was used for survival analysis to detect the effectiveness of these T cell expansion signatures between the experimental and comparator arms. The only difference is that instead of merging the CD8+ and CD4+ T cell expansion signatures, they were kept separate; the patients with an above-median score for either signature (positive-positive, positive-negative, negative-positive) were considered the biomarker-positive group, and the remaining patients were considered the biomarker-negative group. This stratification strategy corresponded to a previous observation that patients in POPLAR who were above-median for either expansion signature had more similar survival patterns with one another than with patients in the negative-negative group. To compare the effectiveness of other signatures between the experimental and comparator arms in IMmotion150 and in POPLAR, the 19-gene dual expansion signature from Wu et al.^13^ was applied, unweighted, as a single signature, while *CD274* (PD-L1), *PDCD1* (PD-1), *CD8A*, *CD4* and *FOXP3* were each treated as its own individual unweighted signature.

## Data availability

All scRNA-seq and scTCR-seq data in this study are available under their original Gene Expression Omnibus (GEO) accession number GSE123814. All bulk RNA-seq data from clinical trials in this study are available under their original GEO accession number GSE139555.

## Code availability

All code used to generate the results will be made freely available from GitHub.

## Acknowledgements

We thank Roshan Kumar, Jennifer Watkins-Yoon, and Joanna Wu for input and reviewing the manuscript, Francisco Adrian for stimulating discussions and support, and HiFiBiO Therapeutics for sponsoring the research.

## Author Contributions

DL performed all primary data analysis and wrote the manuscript. RF provided data interpretation and input on the study design and helped write the manuscript. MM performed the TCR clonotype assessment and helped write the manuscript. LS provided input on the study design and helped write the manuscript. AR conceived the study and wrote the manuscript.

## Supplementary Figures & Tables

**Suppl. Fig. 1.**
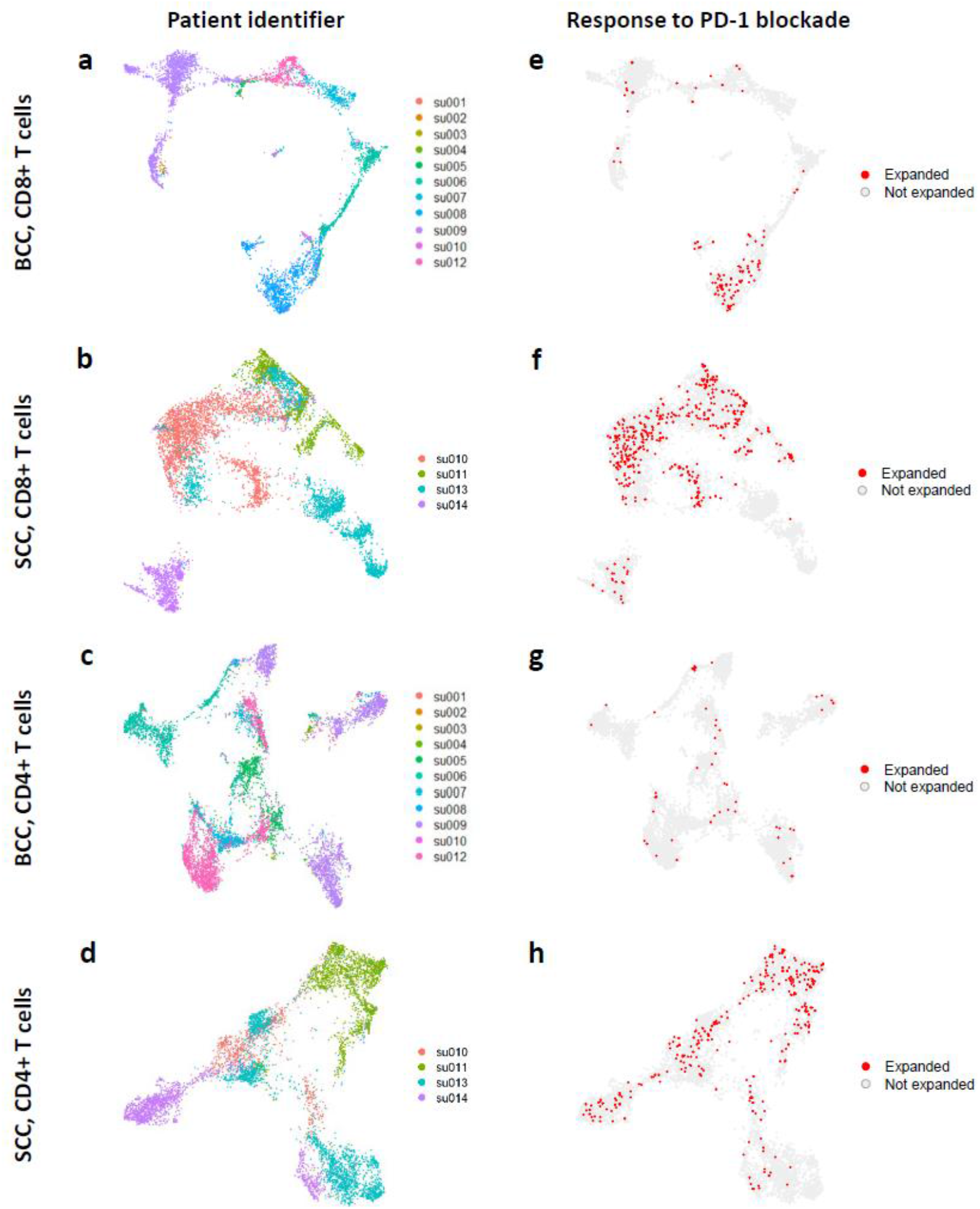
T cells subsets from pretreatment BCC and SCC samples with identification of patient of origin and expansion status. **a-h**, Cells were clustered and visualized with dimension reductions algorithms based on the similarity of their gene expression profiles. Association of single-cells with patient identifiers was indicated (**a-d**). T cells that have expanded in response to anti-PD-L1 treatment have been identified based on scTCR-seq data and are marked in red, all other T cells are displayed in gray (**e-h**).

**Suppl. Fig. 2.**
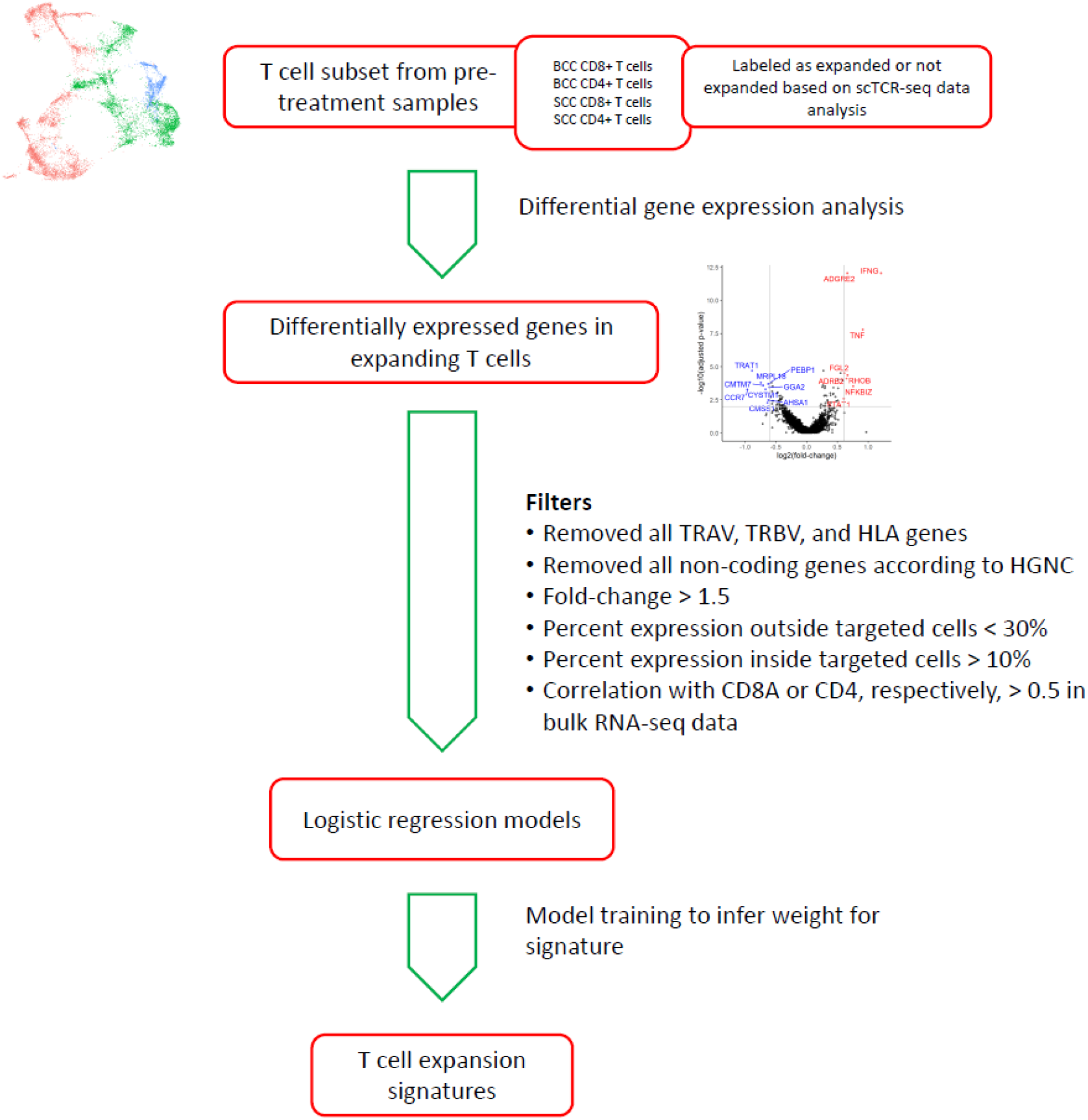
Workflow for predictive gene expression signature inference from single-cell data. Process diagram illustrating the steps in the workflow for predictive gene expression signature inference from scRNA-seq data.

**Suppl. Fig. 3.**
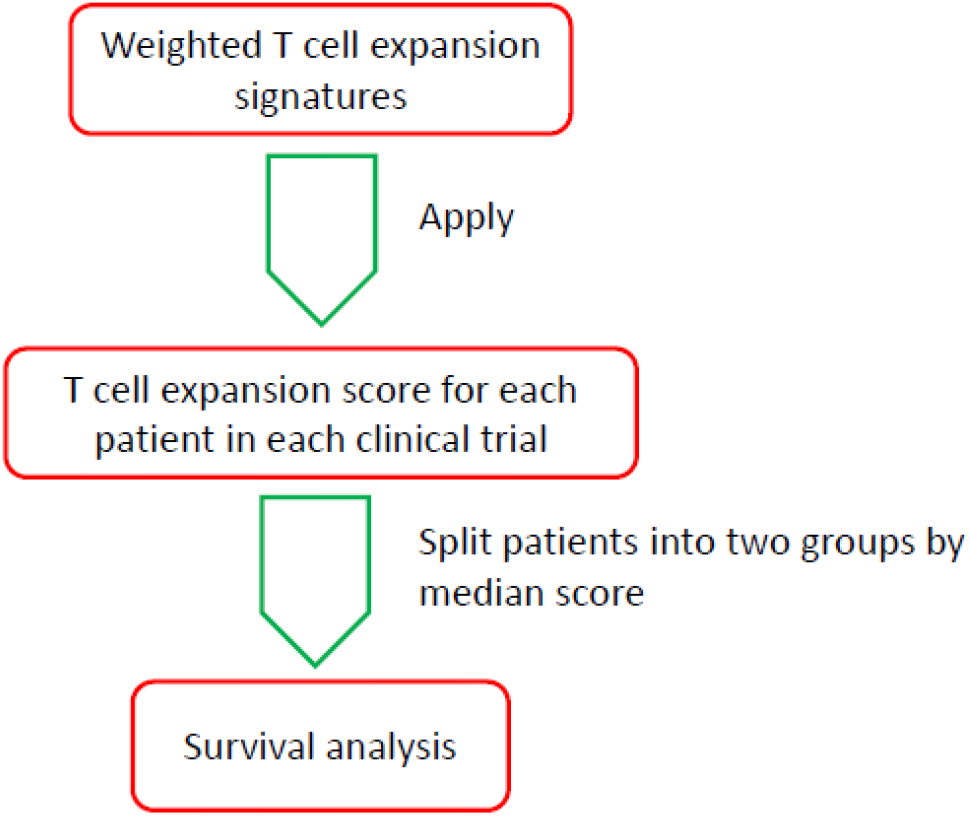
Workflow for application of predictive gene expression signatures to bulk RNA-seq data from clinical trials. Process diagram illustrating the steps in the workflow for application of the predictive gene expression signatures to bulk RNA-seq data from clinical trials.

**Suppl. Fig. 4.**
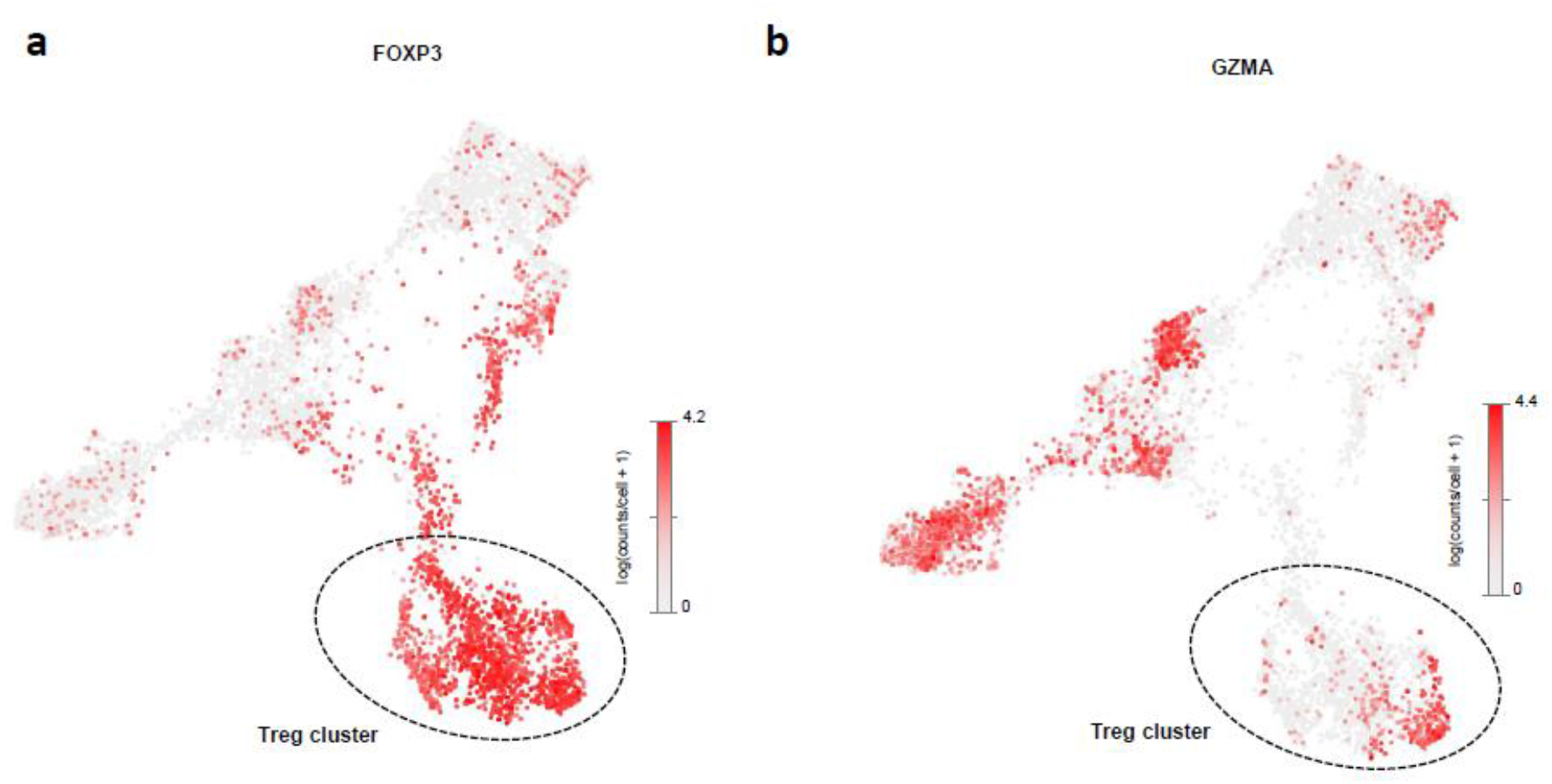
Genes expression of *FOXP3* (a) and *GZMA* (b) contained in the CD4-non-expansion signature identified from CD4+ T cells in SCC patient samples. Strength of expression of each gene is indicated by a color scale. The cluster identified as Tregs is indicate by a dashed circle line.

**Suppl. Tab. 1.** Comparison of hazard-ratios between biomarker positive and negative populations. Hazard ratios and two-sided statistical test from a Cox proportional-hazards model on patients in both groups are shown with significance indicated (* p_val_ < 0.05, ** p_val_ < 0.01, *** p_val_ < 0.001).

| Signature | IMvigor210<br>Atezolizumab | POPLAR<br>Atezolizumab | IMmotion150<br>Atezolizumab | IMmotion150<br>Atezolizumab + Bevacizumab |
| --- | --- | --- | --- | --- |
| **Expansion signature** | **0.65\*\*** | **0.44\*\*** | **0.91** | **0.56\*** |
| Wu et al.<sup>18</sup> | 0.72** | 0.6* | 0.69 | 0.49** |
| *CD274* (PD-L1) | 0.86 | 0.96 | 0.73 | 0.77 |
| *PDCD1* (PD-1) | 0.77* | 0.58* | 0.74 | 0.48** |
| *CD8A* | 0.84 | 0.56* | 0.77 | 0.52** |
| *CD4* | 0.93 | 0.71 | 0.96 | 0.77 |
| *FOXP3* | 0.76* | 0.64* | 0.9 | 0.83 |

